# Loss of Bbs8 leads to cystic kidney disease in mice with reduced acetylation of ciliary alpha-tubulin through HDAC2

**DOI:** 10.1101/2024.03.07.583949

**Authors:** Emilia Kieckhöfer, Julia Günzler, Peter A. Matthiessen, Lena K. Ebert, Christina Klausen, Dagmar Wachten, Metin Cetiner, Max C. Liebau, Thomas Benzing, Helen May-Simera, Bernhard Schermer

## Abstract

Primary cilia dysfunction underlies a group of severe disorders known as ciliopathies. These include Bardet-Biedl syndrome (BBS), which is caused by mutations in BBS genes encoding for components of ciliary protein complexes essential for the assembly and maintenance of primary cilia. As in most ciliopathies, a hallmark feature of BBS is the development of cystic kidney disease. However, the molecular mechanisms linking ciliary dysfunction to cystogenesis remain incompletely understood. Here, we show that Bbs8^-/-^ mice develop late-onset cystic kidney disease accompanied by increased regulated cell death and fibrosis. While the number and length of cilia are not affected, loss of BBS8 reduces K40 acetylation of α-tubulin within primary cilia, compromising ciliary stability. Notably, proteomic analysis revealed a significant upregulation of histone deacetylase HDAC2 in Bbs8^-/-^ kidneys, which we confirmed in Bbs8^-/-^ mouse embryonic fibroblasts (MEFs) and in urine-derived renal epithelial cells (URECs) from a BBS8 patient. We further demonstrate the protein interaction between BBS8 and HDAC2, implicating a disrupted BBS8–HDAC2 regulatory axis in disease pathogenesis. Consistent with a role of excessive HDAC2 activity in the BBS8 deficient cells, pharmacological inhibition of HDAC2 restored tubulin acetylation in BBS8 urine-derived cells. Thus, modulation of HDAC2 activity may represent a strategy to alter ciliary stability in vivo which could explain positive effects of class I specific HDAC inhibitors in models of cystic kidney disease.

## Introduction

The primary cilium is a specialized microtubule-based sensory organelle extending from most mammalian cells’ surfaces (Pazour and Bloodgood, 2008). It consists of the axoneme, ensheathed by the ciliary membrane, the transition zone, and the basal body, which anchors the cilium to the cell body. The transition zone between the basal body and the ciliary shaft is critical for protein trafficking to and from the ciliary compartment (Reiter, Blacque and Leroux, 2012; Garcia-Gonzalo and Reiter, 2017). Tight regulation of the protein composition is required for cilia to function as a hub for chemo- and mechano-transduction, as well as ciliary signaling (Reiter, Blacque and Leroux, 2012). As such, they play a pivotal role in modulating various signaling pathways, e.g. Hedgehog, Wnt, and TGF-β signaling (Fan and Tessier-Lavigne, 1994; Munsterberg et al., 1995; Pourquié et al., 1996). Unlike any other organelle, cilia periodically disappear during the mitotic cell cycle: When cells re-enter mitosis, cilia are typically disassembled to subsequently release the basal body, a modified mother centriole, to serve as microtubule organizing centers at the spindle poles. Disassembly of cilia mechanistically involves the deacetylation of ciliary α-tubulin by HDAC6 and, thus, destabilization of microtubules (Pugacheva et al., 2007; Gopalakrishnan et al., 2023). It is well established that pathogenetic variants in proteins, which alter the structure or function of cilia, lead to a heterogeneous group of genetic diseases and syndromes referred to as ciliopathies (Hildebrandt, Benzing and Katsanis, 2011). Ciliopathies comprise a diverse group of genetic disorders with overlapping phenotypes in different organs and tissues, such as the brain, eye, skeleton, liver, vasculature, and kidney (Forsythe and Beales, 2013; Reiter and Leroux, 2017). Thereby, cystic and fibrotic kidney diseases are a common occurrence among many ciliopathies, leading to the frequent classification of a significant subgroup, namely: renal ciliopathies (Hildebrandt and Zhou, 2007; McConnachie, Stow and Mallett, 2021).

The Bardet-Biedl Syndrome is an archetypical autosomal-recessive ciliopathy characterized by obesity, retinopathy, polydactyly, and kidney disease (Forsythe and Beales, 2013). This syndrome is caused by pathogenic variants of BBS genes encoding for BBS proteins. Most BBS proteins can be classified according to the particular protein complexes which they form. The BBSome complex (Nachury et al., 2007; Loktev et al., 2008) (BBS1, BBS2, BBS4, BBS5, BBS7, BBS8, BBS9, BBS18) is required for trafficking of ciliary membrane proteins whereas, the chaperonin-like complex (BBS6, BBS10, and BBS12), catalyzes the assembly of the BBSome (Seo et al., 2010). The remaining BBS proteins have varying specific functions but are predicted to influence ciliary trafficking. The kidney phenotype in BBS patients is variable with kidneys showing parenchymal, medullary, and most often corticomedullary cysts, but also renal fibrosis, unilateral agenesis, or dysplastic kidneys (Beales et al., 1999; Putoux et al., 2012; Forsythe et al., 2017; Elawad et al., 2022). However, these phenotypes are highly variable among families and could be dependent on mutation type (Putoux et al., 2012; Forsythe et al., 2017).

Studies on various Bbs mutant mice, including Bbs1^M390R^ transgenic, Bbs2^-/-^, Bbs4^-/-^, Bbs5^-/-^, Bbs6^-/-^, Bbs8^-/-^ and Bbs10^-/-^, revealed significant parallels to human disease. These mice exhibit a broad spectrum of vision impairment, obesity, male infertility, and neurological deficits (Kulaga et al., 2004; Mykytyn et al., 2004; Nishimura et al., 2004; Fath et al., 2005; Davis et al., 2007; Rahmouni et al., 2008; Seo et al., 2009; Cognard et al., 2015; Kretschmer et al., 2019; Bentley-Ford et al., 2021; Rödig et al., 2022; Sieckmann et al., 2024). For BBS10, complete knockout resulted in renal abnormalities primarily affecting glomeruli and podocytes without cyst formation, while a tubular epithelial cell-specific knockout did not show any overt phenotype (Cognard et al., 2015). Cyst formation has been observed in Bbs2^-/-^ and Bbs4^-/-^ mice, but again primarily affecting glomeruli (Nishimura et al., 2004; Guo et al., 2011). Loss of Bbs8 has so far shown to cause the most severe retinal phenotype in mice (Tadenev et al., 2011; Dilan et al., 2018; Kretschmer et al., 2019; Schneider et al., 2021), which is not only caused by ciliary defects of photoreceptor cells but also due to ciliary defects in the retinal pigment epithelium (RPE) (Kretschmer et al., 2023; Schneider et al., 2021). BBS8 is one of the key components of the BBSome, which might explain the severity of the Bbs8^-/-^ phenotype. BBS8 (TTC8) is a tetratricopeptide repeat (TPR) protein with a critical role in planar cell polarity and laterality (May-Simera et al., 2010, 2015; Patnaik et al., 2019). Loss of Bbs8 leads to changes in the composition of the octameric BBSome complex, where BBS8 is a direct interactor of the scaffold protein BBS9 (Zhang et al., 2012).

Until now, research into BBS8 function has predominantly focused on embryonic development and ocular health. A subtle renal phenotype, characterized by mild tubular dilation at around 8 months of age, has been reported (Tadenev et al., 2011), but specific roles of BBS8 in maintaining renal cilia and preserving kidney architecture remain largely unexplored. In this study, we report the occurrence of tubular cystic kidney disease in Bbs8-deficient mice at 46 weeks of age, accompanied by renal fibrosis and increased rate of regulated cell death. Interestingly, neither cilia number, nor length differed significantly from wild-type littermates; however, the levels of acetylated tubulin within primary cilia were markedly reduced. We demonstrate that this reduction, which destabilizes cilia, is driven by increased expression of histone deacetylase HDAC2. This effect is confirmed in isolated mouse embryonic fibroblasts (MEFs) and urine-derived renal epithelial cells (URECs) from a BBS8 patient. Thus, beyond potential therapeutic implications of HDAC2 inhibition, our data highlight the importance of not relying solely on acetylated tubulin as a marker for cilia in functional studies, particularly in disease contexts where ciliary stability is affected.

## Methods

### Mouse lines

Bbs8^-/-^ mice have been previously described (Tadenev et al., 2011). All animals were housed, handled, and animal studies conducted, in accordance with approved institutional animal care and use committee procedures. All experiments had ethical approval from the Landesuntersuchungsamt Rheinland-Pfalz and were performed in accordance with institutional guidelines for animal welfare, German animal protection law, and the guidelines given by the ARVO Statement for the Use of Animals in Ophthalmic and Vision Research. Animal maintenance and handling were performed in line with the Federation for Laboratory Animal Science Associations (FELASA) recommendations. Animals were housed in a 12 h light/dark cycle with food and water available ad libitum. For the preparation, mice were weighted, followed by cervical dislocation; blood was collected prior to the perfusion of the kidney with PBS through the aorta. Further, other organs and fat tissue were taken and, as the kidney, processed by either fixation in 4 % formaldehyde and embedded in paraffin or snap-frozen for further tissue analysis. The blood was incubated for 2 h at RT, followed by centrifugation. Serum creatinine levels were measured by the Institute of Clinical Chemistry, University Hospital of Cologne, Germany.

### Immunohistology

For histological analysis, tissue was cut into 2-μm-thick sections and deparaffinized by xylene treatment and rehydration in graded ethanol. For PAS staining, the sections were stained with 0,9 % periodic acid (cat# 3257.1, Roth) and Schiffsches Reagent (cat# 1.09033, Merck) both for 10 min embedded into washing steps with H_2_O. Finally, to visualize nuclei in blue, the samples were stained with Mayer’s Haematoxylin for 20 s. For the Masson staining the Masson-Goldner’s trichrome staining kit (cat# 3459, Roth) was used and performed according to the manufacturer’s instructions. After dehydration of the sections, they were embedded with Histomount (HS-103, National Diagnostics).

### Cyst index analysis

The cyst index was calculated for whole slide images using open-source software for bioimage analysis QuPath (v0.4.0) (Bankhead et al., 2017). Cysts were detected using an Artificial Neural Network-based pixel classifier. Initially detected cysts were filtered for the minimal area of 400 µm² and a minimum circularity value of 0.35. Plots were generated using the Plots of Data web app (Postma and Goedhartid, 2019). For statistical analysis, all results were normalized to the control followed by a two-tailed Student’s t-test (p<0.05).

### Proteome and phosphoproteome analysis

For each biological replicate a quarter of kidney tissue was used and dounced with a Wheaton Dounce tissue grinder in urea buffer (8 M Urea, 50 mM ammonium bicarbonate) supplemented with Halt protease-phosphatase-inhibitor cocktail (Thermo Scientific™). After clearing of the sample (16.000 xg, 1 h at 4°C), the lysates were reduced (10 mM dithiothreitol), alkylated (50 mM chloroacetamide) and digested (LysC; 1:75). Samples (800 µg) were diluted to 2 M urea and subjected to tryptic digestion (1:50). After overnight incubation, phosphoenrichment was performed in the CECAD proteomics facility using the Thermo Scientific™ Kit High Select TiO2 Kit (#A32993). All samples were analyzed as well by the CECAD proteomics facility on a Q Exactive Plus Orbitrap mass spectrometer that was coupled to an EASY nLC (both Thermo Scientific™). Peptides were loaded with solvent A (0.1% formic acid in water) onto an in-house packed analytical column (50 cm, 75 µm inner diameter, filled with 2.7 µm Poroshell EC120 C18, Agilent). Peptides were chromatographically separated at a constant flow rate of 250 nL/min using the following gradient: 3-5% solvent B (0.1% formic acid in 80 % acetonitrile) within 1.0 min, 5-30% solvent B within 121.0 min, 30-40% solvent B within 19.0 min, 40-95% solvent B within 1.0 min, followed by washing and column equilibration. The mass spectrometer was operated in data-dependent acquisition mode. The MS1 survey scan was acquired from 300-1750 m/z at a resolution of 70,000. The top 10 most abundant peptides were isolated within a 1.8 Th window and subjected to HCD fragmentation at a normalized collision energy of 27%. The AGC target was set to 5e5 charges, allowing a maximum injection time of 55 ms. Product ions were detected in the Orbitrap at a resolution of 17,500. Precursors were dynamically excluded for 25.0 s. All mass spectrometric raw data were processed with MaxQuant (Tyanova, Temu and Cox, 2016) (version 2.2.0.0) using default parameters against the UniProt canonical murine database (UP10090, downloaded 20.01.2023) with the match-between-runs option enabled between replicates. Samples were sorted into two parameter groups, either containing the enriched or non-enriched samples. Enriched samples had the phosphorylation (STY) variable modification added, whereas non-enriched samples were quantified by LFQ. A follow-up analysis was done in Perseus 1.6.15 (Tyanova et al., 2016). Results were cleaned up by removing hits from the decoy database, the contaminant list and, in case of non-enriched fractions, those only identified by modified peptides were removed. Afterwards, results were filtered for data completeness in at least one condition and LFQ values (WP) or intensities (PP), imputed using sigma downshift with standard settings. Finally, FDR-controlled two-sided t-tests between sample groups were performed (S0=0, FDR≤0.05) as well as a 1D annotation enrichment using Perseus (version 1.6.15.0).

### Isolation and immortalization of mouse embryonic fibroblasts (MEFs)

For the isolation of mouse embryonic fibroblasts (MEFs), timed matings were set up with one male and two females of the desired genotype. At day 13, the pregnant mouse was anesthetized using isoflurane (Piramal Healthcare) followed by a cervical dislocation. The lower abdomen was opened by an abdominal incision to extract the two uterine horns. Embryos were isolated and transferred into a 24-well plate filled with PBS. The head and the red organs (heart and liver) were removed. The rest of the embryo was placed into a 12-well plate filled with 2 ml ice-cold 0.25 % Trypsin/PBS (diluted from 2.5 % Trypsin, Gibco). The embryos were chopped into small pieces and incubated overnight at 4 °C. The trypsin solution was discarded and the remaining Trypsin/tissue mixture was incubated for 30 min in a 37 °C water bath. Afterward, medium (composition: DMEM/Glutamax, 10 % FCS, 1 % sodium pyruvate (100x), 1 % Pen Strep), was added, and the cell suspension was pipetted vigorously up and down to break up the digested tissue into a single cell suspension. After 1 min, the cell suspension was collected, to remove the sedimentation of the remaining tissue. This step was repeated and the cell suspension was filtered through a 100 μm cell-strainer (Corning). Cells were plated and after 24 h, the medium was changed. Immortalization of MEFs was performed as described previously (Todaro and Green, 1963). Briefly, cells are split every three days and seeded with the same cell density. From passage three onwards, cells were seeded on at least two 10 cm culture dishes. After around 15 passages, cells started to regrow. When MEFs were immortalized, frozen back-ups were made.

### Isolation, immortalization, and BBS8 re-expression of urine renal epithelial cells (URECS)

BBS8^c.915delG^ patient urine (male; age 14) was collected at the University Hospital Essen, Germany, in cooperation with the Network for early onset cystic kidney disease (Neocyst). Urine sample was centrifuged (10 min, 400 x g), and the pellet was resuspended in 10 ml washing buffer (sterile 1x PBS, 100 U Penicillin/Streptomycin (Gibco™), 500 ng/ml Amphotericin B (Sigma)). After centrifugation (10 min, 200 x g) the supernatant was removed leaving ∼0.2 – 0.5 ml in the tube. The pellet was resuspended in primary medium (PUC; DMEM-F12 (Sigma), REGM SingleQuot kit (Lonza, cat# CC-4127), 10% FBS (Gibco™), 21mM GlutaMAX (Gibco™), 1.0% Penicillin/ Streptomycin (Gibco™), 500 ng/ml Amphotericin B (Sigma)) and plated in a single well of a galantine-coated 12-well plate. The plate was incubated for 48 h at 37 °C, before the first PUC medium change. With the first visible colonies, PUC medium was changed to normal growth medium (DMEM-F12 (Sigma), 10% FBS (Gibco™), 21mM GlutaMAX (Gibco™), 1.0% Penicillin/ Streptomycin (Gibco™)). Reaching a confluency of 60-80 %, cells were split in two wells of a 6-well plate using TrypLE Select Express (Gibco™). One well was expanded, and stored as primary cells, and one well was used for immortalization. Immortalization was started by adding 1 ml of previously produced virus supplemented with 1 µl of Polybrene 8mg/ml (Sigma) to 40 % confluent cells and incubated for 24 h at 37°C. The virus was removed and replaced by normal growth medium. Cells were split thrice, while expansion in multiple 10 cm culture dishes before frozen back-ups were made. Immortalized urine-derived renal epithelial cells (URECS) were additionally expressed exogenous with viruses produced with pLenti6.3 F.hBBS8 and pLenti6.3 F.GFP in HEK293T cells. Sterilized harvest virus supplemented with Polybrene was added to 40 % confluent cells and incubated for 24 h at 37°C. After removing the virus, the cells were selected by adding normal growth medium supplemented with 0,001 mg/ml Blasticidin (Invivogen). Cells were split three times before selection was stopped and cells were used for experiments.

### Cell culture and treatment

Human embryonic kidney 293T cells (HEK293T, ATCC®), mIMCD3 cells, MEFs and URECs were cultivated at 37°C and 5% CO_2_. Thereby, HEK293T cells were maintained in DMEM + GlutaMAX™ medium (Gibco™) supplemented with 10% fetal bovine serum (FBS, Gibco™), MEFs in DMEM/F-12 + GlutaMAX™ medium (Gibco™) complemented with 10% FBS and 1% penicillin-streptomycin.URECs and mIMCD3 cells were cultured in normal growth medium (DMEM-F12 (Sigma), 10% FBS (Gibco™), 21mM GlutaMAX (Gibco™). For URECs, 1.0% Penicillin/ Streptomycin (Gibco™) was added. All cell lines were tested negative for mycoplasma (PCR Mycoplasma Test Kit I/C, PromoKine). To induce a higher amount of ciliogenesis, mIMCD3 cells were grown in serum-free media for 24 h at 37°C. HEK293T cells were transiently transfected with5 µg (IP) or 6 µg (Interactome) of F.hBBS8 in pcDNA6 using the calcium phosphate method. As negative control, equal amounts of F.EPS^1-225^ in pcDNA6 were used. The HDAC2 inhibitor Tucidinostat (TUC, 160 nM; Biozol, cat# HY-109015) treatment was performed in serum-free media at 37°C for 24 h. Control treatment was performed with DMSO (AppliChem). For IF stainings, confluent mIMCD3 cells were transfected with 5 µg of either F.EPS or F.HDAC2 using Lipofectamine®3000 (Invitrogen) in starvation medium and fixed 24 h after transfection. Every plasmid used for this study added an N-terminal F9-tag to the protein of interest, which refers to a FLAG-tag followed by 9 histidine residues. Only mHDAC2 was expressed with a single FLAG-tag.

### Harvest and fixation

Cells were seeded in dishes or on glass coverslips in 12 well plates and incubated for 24 h, prior to starvation or TUC treatment; both for an additional 24 h at 37°C. To harvest cells, they were washed with PBS (1x), scraped in fresh PBS (1x), and centrifuged for 10 min, 10.000 rpm at 4°C. Cell pellets could be used for western blot or RNA isolation. Harvest of whole cell lysate samples was performed by resuspending the cells in cold 1x PBS followed by centrifugation for 5 min, 4.000 rpm at 4°C. Afterwards cells were resuspended in 1x Laemmli with DTT and boiled for 5 min at 95°C. For Immunofluorescence staining cells were fixed by adding 4% PFA (Walter CMP GmBH & Co) for 5 min, followed by methanol (Roth) for 4 min. Fixed cells were washed thrice with PBS (1x) before staining.

### Cilia stability assay

Depolymerization of microtubules was induced by incubating starved cells for 30 min on ice. The subsequent fixation was continuously performed on ice. The untreated control cells were fixed at room temperature. Fixed cells were stored at 4°C in PBS until staining and quantifying cilia.

### Quantitative real-time PCR

RNA isolation was performed from kidney tissue samples, thereby using one-quarter of a kidney. Tissue was ground with BeadBeater (Roth) using a Precelly24 with 5.000 rpm two times for 30 s in Tri-Reagent. RNA isolation from cells was performed by resuspending snap-frozen cell pellets in Phenol. RNA extraction was performed with the Direct-zol RNA Miniprep kit (Zymo Research) following the manufacturer’s instructions, including a DNase1 treatment step. Before the reverse transcription using the High-Capacity cDNA Reverse Transcription kit (Applied Biosystems), RNA concentration and sample quality were assessed on a Nanodrop spectrophotometer (Peqlab). mRNA was assessed by qPCR with SYBR Green (Thermo Scientific™) using mHprt1 or hHPRT1 as endogenous control. The qPCR experiments were performed on a QuantStudio Q5 Real-time PCR System (ThermoFisher Scientific). For data analysis, all results were normalized to the housekeeping gene Hprt1 using the delta-delta CT followed by a two-tailed Student’s t-test (p<0.05).

### Co-Immunoprecipitation

Co-immunoprecipitation (Co-IP) was performed as previously described (Habbig et al., 2011) using the IP Buffer (20 mM Tris, 1% (v/v) TritonX-100, 50 mM NaCl, 15 mM Na_4_P_2_O_7_, 50 mM NaF, pH 7.5) supplemented with inhibitors (44 μg/μl PMSF, 2 mM Na_3_VO_4_) in four replicates. Input samples (lysates) were collected, and the remaining samples were incubated with α-FLAG M2 Beads (A2220, Sigma-Aldrich) for 2 h. Samples were denatured and reduced in 2x Laemmli before SDS-PAGE. For proteomic analysis, the IP Buffer was supplemented with cOmplete™ Protease Inhibitor Cocktail (Roche). After sonication, the lysates were first centrifuged 30 min at 14.000 g at 4°C followed by ultracentrifugation for 45 min at 100.000 g and 4°C. After three washing steps the beads were incubated with 5% SDS in PBS for 5 min at 95 °C, followed by reduction with 10 mM dithiothreitol and alkylation with 40 mM chloroacetamide.

### Interactome

Samples were analyzed by the CECAD Proteomics Facility on an Orbitrap Exploris 480 (Thermo Scientific, granted by the German Research Foundation under INST 1856/71-1 FUGG) mass spectrometer equipped with a FAIMSpro differential ion mobility device that was coupled to an Vanquish neo in trap-and-elute setup (Thermo Scientific). Samples were loaded onto a precolumn (Acclaim 5μm PepMap 300 μ Cartridge) with a flow of 60 μl/min before reverse-flushed onto an in-house packed analytical column (30 cm length, 75 μm inner diameter, filled with 2.7 μm Poroshell EC120 C18, Agilent). Peptides were chromatographically separated with an initial flow rate of 400 nL/min and the following gradient: initial 2% B (0.1% formic acid in 80 % acetonitrile), up to 6 % in 3 min. Then, flow was reduced to 300 nl/min and B increased to 20% B in 26 min, up to 35% B within 15 min and up to 98% solvent B within 1.0 min while again increasing the flow to 400 nl/min, followed by column wash with 98% solvent B and re-equilibration to initial condition. The FAIMS pro was operated at −50V compensation voltage and electrode temperatures of 99.5 °C for the inner and 85 °C for the outer electrode. The mass spectrometer was operated in data-dependent acquisition top 24 mode with MS1 scans acquired from 350 m/z to 1400 m/z at 60k resolution and an AGC target of 300%. MS2 scans were acquired at 15 k resolution with a maximum injection time of 22 ms and an AGC target of 300% in a 1.4 Th window and a fixed first mass of 110 m/z. All MS1 scans were stored as profile, all MS2 scans as centroid. All mass spectrometric raw data were processed with Maxquant (version 2.4) (Tyanova, Temu and Cox, 2016) using default parameters against the Uniprot HUMAN canonical database (UP5640) with the match-between-runs option enabled between replicates. Follow-up analysis was done in Perseus 1.6.15 (Tyanova et al., 2016). Protein groups were filtered for potential contaminants and insecure identifications. Remaining IDs were filtered for data completeness in at least one group and missing values imputed by sigma downshift (0.3 σ width, 1.8 σ downshift). Afterwards, FDR-controlled two-sided t-tests were performed (S0=0, FDR≤0.05). Protein structure prediction was done using the Alphafold server (Abramson et al., 2024), while ChimeraX has been used for visualization (Pettersen et al., 2021; Meng et al., 2023).

### Immunoblotting

Confluent grown, and starved MEF cells were harvest with RIPA buffer (50 mM Tris/HCl, 150 mM NaCl, 1% (v/v) NP-4O, 0.5% (w/v) Sodium deoxycholate, 0.1% (w/v) SDS), supplemented with 1% Halt Protease and Phosphatase Inhibitor Cocktail (Thermo Scientific™) (Brücker et al., 2023). Cells were lysed on ice and sonicated for 2 s. Protein concentration of lysates was determined and loaded onto a 10% SDS-PAGE and subsequently transferred to a PVDF-FL membrane (Millipore). URECSs cell pellets were lysed in adjusted inhibitor-supplemented RIPA buffer (50 mM Tris/HCL pH 7.5, 150 mM NaCl, 0.1 % NP-40, 0.5 % Na-Deoxycholat, 0.1 % SDS) supplemented with Benzonase® (70746-3 Millipore), cOmplete^TM^ (4693159001, Roche) and PhosSTOP^TM^ (4906845001, Roche), on ice for 30 min. This is followed by centrifugation for 1 h, 14.000 rpm at 4°C, determination of protein concentration, and supplementing and boiling with 5x Laemmli. Samples were loaded onto a gel as before and transferred to a PVDF membrane (Millipore), including whole cell lysates of mIMCD3 cells in 1x Laemmli. After blocking, membranes were incubated overnight at 4°C in primary antibodies (FLAG, F7425, Sigma-Aldrich, 1:1000; HDAC2, ab32117 Abcam, 1:1000; acetylated Tubulin, T6793 Sigma, 1:1000; Calnexin, 10427-2 AP ProteinTech, 1:1000). Secondary antibodies were incubated for 1 h at RT either fluorescent (Li-COR Biosciences IRDye680 and IRDye800 1:10.000: rb680, 925-68073; mm680, 925-68072; rb800, 925-32213) or horseradish peroxidase-conjugated antibodies. Fluorescent signals were visualized using the Odyssey Infrared Imaging System 2800 (Li-COR), for peroxidase-conjugated ECL detection, and images were acquired with a Fusion Solo S (Vilber Lourmat Germany GmbH, Eberhardzell, Germany).

### Immunofluorescence, Immunochemistry and TUNEL staining

The kidney tissue staining of 4 µm (cilia staining) and 2 µm (other stainings) fixed sections were performed as previously described (Dafinger et al., 2021). The primary antibodies (NF-kB, #8242 Cell Signaling, 1:1000; CD3, MCA-1477 Biorad, 1:100; acetylated Tubulin, T6793 Sigma, 1:1000; Slc12a3, HPA028748, Sigma Aldrich, 1:500; ARL13B, 17711-1-AP ProteinTech, 1:500) were incubated overnight at 4°C, followed by incubation with secondary antibodies. For immunochemistry, the DBA kit (K3468 DAKO, 30 min for NF-kB; SK-4105 Vector, 5 min for CD3) was used and samples counterstained with Hematoxylin. For immunofluorescence, fluorophore-coupled antibodies (Jackson ImmunoResearch, 1:500: anti-mouse-Cy5, # 715-175-150; anti-rabbit A647, 711-605-152; anti-mouse-Cy3, 715-165-150; and FITC-Lotus Tetragonolobus Lectin (LTL), FL-1321-2; Vector laboratories, 1:500) were used for 1 h at RT. Samples were mounted after a short incubation with Hoechst 33342 (ThermoFisher Scientific, 1:5000) with ProLong™ Diamond (ThermoFisher Scientific). The DeadEnd™ Fluorometric TUNEL System (Promega) was performed following the manufacturer’s instructions, with the exception that the samples were mounted, with a pre-incubation of Hoechst with ProLong™ Diamond. Fixed MEFs were quenched with 50 mM NH_4_Cl for 10 min, before permeabilisation with 0,3% PBS-TritonX-100 for 20 min. Antibodies were diluted in Fish-Block (0.1 % (w/v) ovalbumin, 0.5 % (w/v) fish gelatine, in PBS), supplemented with 0,3% TritonX-100. The primary antibody (ARL13B 1:800 cat# ab136648, Abcam; acetylated Tubulin, 1:800 cat# T6793, Sigma) were incubated overnight at 4°C followed by the secondary antibodies (anti-rabbit 488, A11034 Invitrogen, 1:400; anti-mouse 555, A31570, Invitrogen, 1:400; DAPI, Roth, 1:400). Finally, coverslips were mounted with Fluoromount-G (ThermoFisher, 00-4958-02) and imaged with the Leica microscope CTR6000, with DM6000B Laser and DFC360FX camera. MEF cell images were deconvoluted with the Leica imaging software LASX. Cilia number and length were determined with the open-source Fiji software (Schindelin et al., 2012). Fixed BBS8^c.915delG^ URECs were blocked for 1 h in PBS containing 0.1 % triton-X-100 and 5 % normal donkey serum. Fixed mIMCD3 cells were permeabilized for 10 min with 0.1 % triton X-100 diluted in PBS followed by a 1 h blocking step with 10 % NDS in PBS supplement with Ca²^+^ and Mg²^+^. The primary antibody (ARL13B 1:1000 cat# ab136648, Abcam; ARL13b 1:400, cat# 17711-1-AP, Proteintech; phospho-HDAC6 1:500 cat# PA5-105035, Invitrogen; acetylated Tubulin 1:1000, Sigma; beta-Tubulin 1:50, cat# E7, DSHB) were incubated at 4°C. Secondary Antibodies (donkey anti-mouse Cy3, cat# 715-165-150, Jackson ImmunoResearch; donkey anti-rabbit A647, cat# 711-605-152, Jackson ImmunoResearch; DYKDDDDK Tag (L5), Alexa Fluor™ 488, 1:500; Invitrogen) were incubated at RT. Nuclei staining was performed with Hoechst 33342 Solution (1:5000, cat# H3570 Invitrogen). Finally, coverslips were mounted with ProLong™ Diamond w/o DAPI (cat# P36965, Thermo Fisher Scientific). URECs and IMCD3 cells were imaged with the AxioObserver microscope with an axioCam ICc1, Axiocam 702 mono, Apoptome system (Carl Zeiss MicroImaging, Jena, Germany). Images were exported through Zen Blue Software by Zeiss and aligned together using the free design tool Inkscape.

### Quantification and statistical analysis

Data are expressed as mean1±1standard deviation (SD). All experiments were performed in at least 3 independent biological replicates. The data were statistically analyzed with GraphPad Prism version 9.5.1 unless otherwise mentioned. Statistical testing was performed as indicated in the figure legends.

## Results and discussion

### Cystic kidney disease in Bbs8 knockout mice is accompanied by inflammation and fibrosis

To investigate the function of BBS8 in the kidney, we analyzed the kidneys of a conventional Bbs8^-/-^ mouse line (Tadenev et al., 2011) where a mild renal phenotype with moderately dilated tubules in the deep cortical region has previously been reported at 8 months of age. Consistent with this, no apparent renal abnormalities were observed at 24 weeks. Therefore, we extended the observation period to 46 weeks. At this age, Bbs8^-/-^ mice exhibited marked obesity, with a body weight approximately 30% higher than that of control animals (Fig. 1A), consistent with a typical BBS phenotype. In addition, signs of renal involvement began to emerge, as indicated by a trend toward elevated serum urea levels (Fig. 1B) and a slight increase in kidney weight (Supp. Fig. 1A), although these changes did not reach statistical significance. PAS staining of kidney sections revealed cyst formation and tubular dilatation (Fig. 1C), with a cyst index approximately six-fold higher in Bbs8^-/-^ compared to control mice (Fig. 1D). Further analysis revealed that although the size of individual cysts detected in Bbs8^-/-^ was similar to the size of dilated tubules and cysts occasionally found in age-matched control animals (Supp. Fig. 1B), the number of dilated tubules and cysts was significantly increased (Supp. Fig. 1C). Staining for proximal and distal tubules in the kidney revealed that the majority of cysts originated from distal tubules, as they were positive for Slc12a3 (Fig. 1E). Furthermore, Bbs8^-/-^ kidneys showed a significant increase in inflammation and fibrosis in PAS and Masson’s trichrome stained sections (Fig. 1F). In line with inflammation and fibrosis, we observed an accumulation of T-cells (CD3^+^), as well as an increased nuclear expression of the NF-кB subunit RelA/p65, in particular in inflammation-rich areas (arrowheads) but also in kidney tubules (arrows), in Bbs8^-/-^ kidney tissue (Fig. 1G). Increased TUNEL-positive cells indicated ongoing cell death in Bbs8^-/-^ kidneys (Fig. 1H). To unravel the role of potential players involved in these processes of cell death, we found the mRNA transcription of RelA/p65, as well as, of the pyroptosis markers Nlrp3 and GsdmD to be significantly increased (Fig. 1I). Taken together, Bbs8^-/-^ mice, in addition to obesity, fatty liver disease, retinal degeneration, and occasional polydactyly, develop a relatively late-onset cystic fibrotic kidney disease accurately reflecting the patient renal manifestation. Cyst formation predominantly originates from the distal tubular segments and increased renal expression of genes associated with pyroptosis and the inflammasome, key factors of renal fibrosis and inflammation.

**Figure 1.**
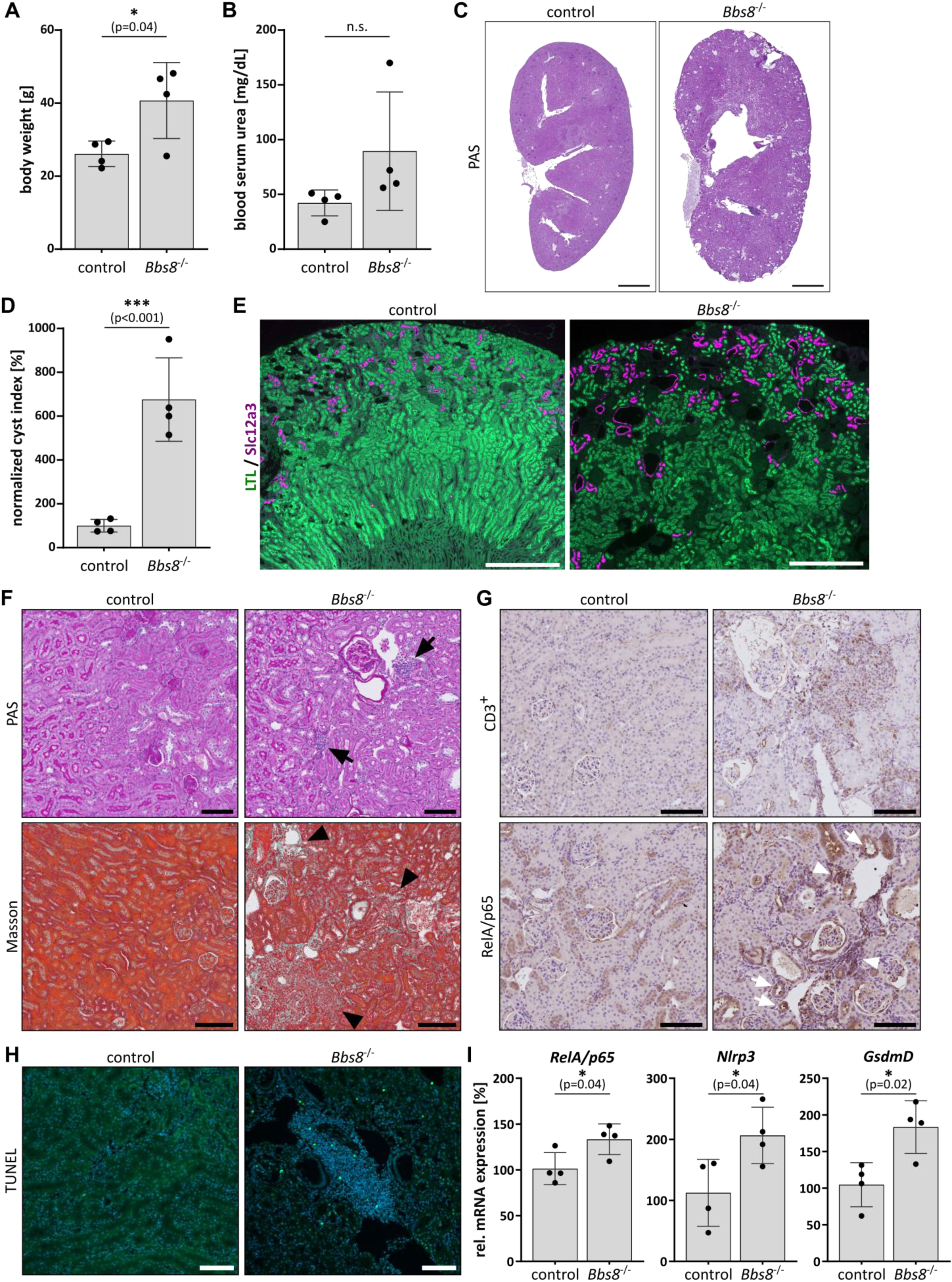
Bbs8^-/-^ mice develop late-onset cystic kidney disease with fibrosis and inflammation. 46-week-old Bbs8^-/-^ compared to control (Bbs8^+/+^) mice showed a significantly increased (A) body weight (n=4) and an increasing trend (B) of measured urea level of blood serum (n=4). (C) PAS staining of the whole kidney in control and Bbs8^-/-^ animals, showing cyst formation in the knockout; Scale bar: 1 mm. (D) Cyst index described in a cyst-to-tissue area ratio, normalized to control animals (n=4). (E) Staining of distal (Slc12a3, magenta) and proximal tubules (LTL, green) in the kidney showing cyst formation in the distal convoluted tubules; Scale bar: 500 µm. (F) Representative microscopic images of the PAS staining (higher magnified area), showing clusters of inflammation (Arrow; Scale bar: 200 µm). The additional Masson’s trichrome staining, displayed fibrosis in Bbs8^-/-^ animals (Arrowhead; Scale bar: 300 µm). (G) Representative microscopic images of CD3^+^ and NF-kB (RelA/p65) expression revealed inflammation in Bbs8^-/-^ animals; Scale bar: 300 µm. (H) TUNEL staining of control and Bbs8^-/-^ kidney tissue; Scale bar: 100 µm. (I) Quantitative real-time PCR of RelA/p65 and inflammasome genes in control and Bbs8^-/-^ kidney samples (n=4). All of the statistical analyses were performed using a two-sided Student’s t-test (p-value: <0.001***; 0.002**; 0.033*; ns1=10.12).

### The loss of Bbs8 affects ciliary tubulin acetylation and primary cilia stability

Since BBS8 is a component of the BBSome and the role of cilia in Bardet-Biedl syndrome (BBS) is well established, we next examined primary cilia in kidney tissue from Bbs8^-/-^ mice. In contrast to mouse olfactory tissue, where a dramatic loss of cilia had been described (Tadenev et al., 2011), we did not observe any significant difference in the number and length of cilia in kidney sections of Bbs8^-/-^ mice compared to wildtype littermates (Fig. 2A). Since quantification of cilia in tissue sections has technical limitations, we also used mouse embryonic fibroblasts (MEFs) generated from Bbs8^-/-^ and control mice. A high percent of MEFs typically form cilia, and quantification of cilia number and length via ARL13B staining revealed no significant difference in serum-starved MEFs (Fig. 2B), confirming our finding in the kidney sections. As an additional model, we isolated URECs from a BBS8 patient with a homozygous BBS8 variant (BBS8^c.915delG^) that results in a frameshift and premature protein termination (p.Met305Ilefs*15). In these cells we re-expressed F.hBBS8 or F.GFP via lentiviral gene transfer. Expression was validated by immunoblotting (Suppl Fig. 2A). Consistently, cilia numbers as determined by ARL13B staining were not significantly different in these cell lines (Fig. 2C). Surprisingly, however, we observed a discrepancy between ARL13B staining and acetylated tubulin, another classical ciliary marker: in kidney sections (Fig. 2E), as well as in MEFs (Fig. 2F) and URECs (Fig. 2G), a significant number of cilia in the BBS8-deficient group identified by ARL13B staining were negative for acetylated tubulin staining. Taking into account that tubulin K40 acetylation is important for microtubule and, thus, ciliary stability, we analyzed this in the BBS8^c.915delG^ URECs with and without re-expression of F.hBBS8 by incubating the cells on ice to depolymerize microtubules and destabilize cilia. This assay revealed that upon treatment, the amount of cilia was significantly reduced in BBS8^c.915delG^ F.GFP URECs as compared to the rescued cells (Fig. 2H/I).

**Figure 2.**
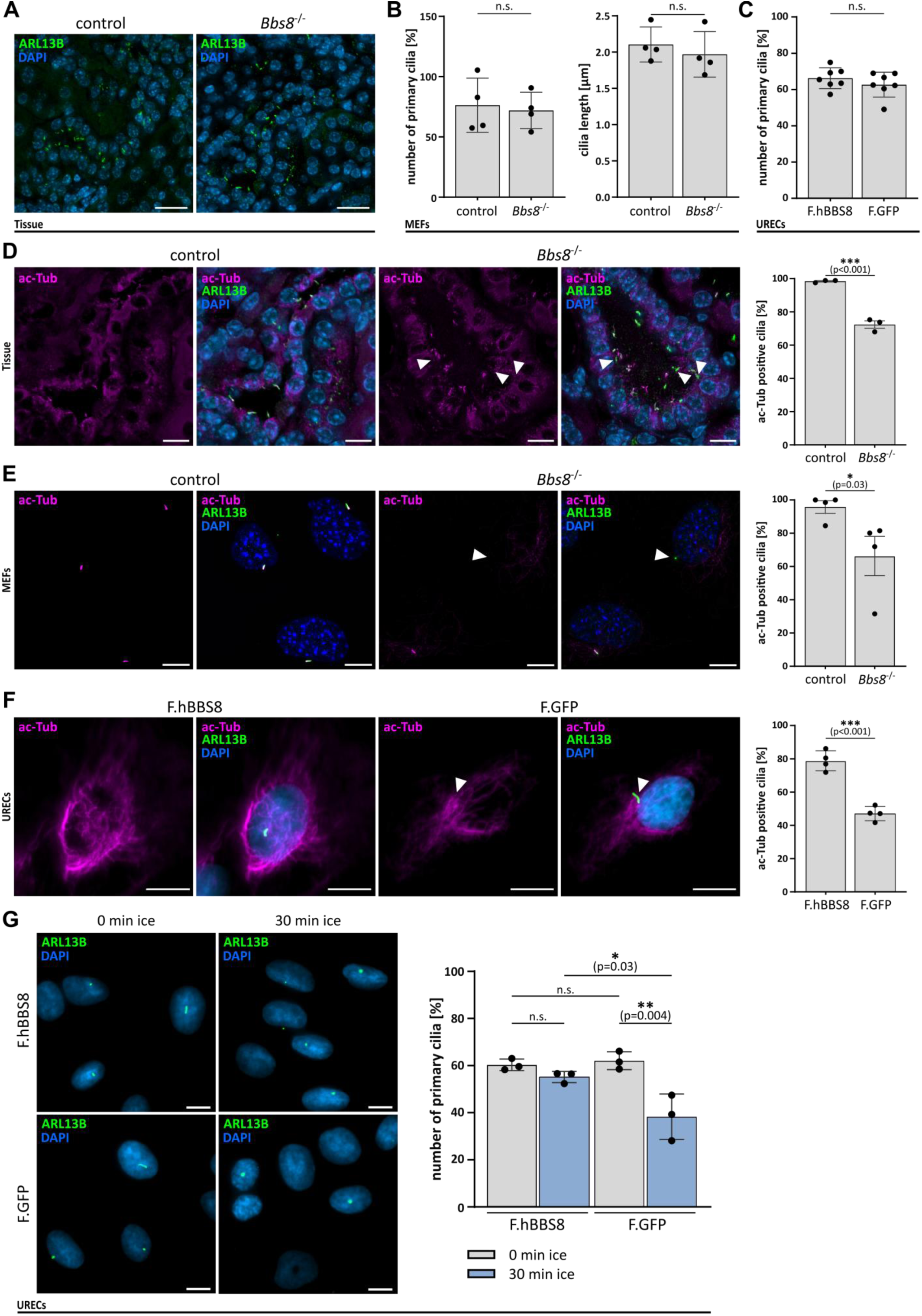
BBS8 deficiency results in reduced ciliary tubulin acetylation and stability without significantly affecting the number or length of cilia. (A) Control and Bbs8^-/-^ kidney tissue stained for cilia using ARL13B antiserum (Scale bar: 50 µm). (B) Bbs8^-/-^ and control mouse embryonic fibroblasts (MEF) quantified for the number and length of cilia by ARL13B staining (n=4), statistically analyzed using an unpaired Student’s t-test (p-value: <0.001***; 0.002**; 0.033*; ns1=10.12). (C) BBS8^c.915delG^ and BBS8 re-expressing URECs stained for cilia using ARL13B antiserum (n=7), statistically analyzed using an unpaired Student’s t-test (p-value: <0.001***; 0.002**; 0.033*; ns1=10.12). (D-F) Representative images of primary cilia co-stained for ARL13B (green) and acetylated-tubulin (ac-Tub, magenta; scale bar: 10 µm); graphs showing the percentage of cells with a ac-Tub positive cilium from the total amount of cilia, as identified by ARL13B staining. Arrowhead indicates ac-Tub negative cilia. Statistically analyzed using an unpaired Student’s t-test (p-value: <0.001***; 0.002**; 0.033*; ns1=10.12). Staining and analysis were performed in (D) control and Bbs8^-/-^ kidney tissue (n=3), (E) cultured Bbs8^-/-^ and control MEFs (n=4), (F) BBS8^c.915delG^ patient-isolated URECs stably expressing either F.hBBS8 or F9.GFP (n=4). (G) Cilia stabilization assay was performed in BBS8^c.915delG^ patient-isolated URECs stably expressing either F.hBBS8 or F.GFP. Destabilization was induced by keeping the cells for 30 min on ice. Representative images of primary cilia stained for ARL13B (green; n=3; Scale bar: 10 µm). Statistical analysis was performed by using a one-way ANOVA followed by a two-sided Student’s t-test (p-value: >0.001***; 0.002**; 0.033*; ns1=10.12).

Thus, the loss of BBS8 does not affect cilia number but leads to a reduction of acetylated tubulin within the cilium, which in turn impacts ciliary stability.

### Proteomic analysis of Bbs8-deficient kidneys

To gain more insights into the kidney pathology and to decipher the mechanism contributing to the loss of tubulin acetylation, we performed unbiased proteomic expression analysis of kidney lysates from Bbs8^-/-^ and control mice. The principal component analysis clearly separated the two genotypes in both datasets (Fig. 3A). After quality control (QC), we found a total of 2926 proteins in the whole proteome (WP) (Fig. 3B; Suppl. Table 1). Based on the Student’s t-test (S0=0; FDR≤0.05), only 10 proteins were significantly up- and 16 significantly downregulated in Bbs8^-/-^ kidney lysates compared to control (Fig. 3C). Among those differentially expressed proteins were several proteins previously associated with ciliopathies, cystogenesis, and kidney disease progression: For example, dynein cytoplasmic 2 heavy chain 1 (DYNC2H1) and dystrobrevin binding protein 1 (DTNBP1) were found to be downregulated in the knockout. DYNC2H1, a subunit of the IFT-dynein motor, drives retrograde IFT-rafts and plays a role in the formation of the ciliary transition zone. Mutations lead to a skeletal ciliopathies, namely Jeune asphyxiating thoracic dystrophy, short-rib polydactyly, and also to non-syndromic retinal degeneration (Jensen et al., 2018; Vig et al., 2020; Hammarsjö et al., 2021; Piceci-Sparascio et al., 2023). Mutations in DTNBP1, encoding a subunit of the biogenesis of lysosome-related organelles complex-1 (BLOC-1), cause late-onset cystic kidney disease in mice (Monis, Faundez and Pazour, 2017). Proteins with increased expression compared to control included retinol-binding protein 4 (RBP4), uromodulin (UMOD), nucleoporin 98 (NUP98), and histone deacetylase 2 (HDAC2), each of which has previously been associated with kidney diseases or primary cilia (Zaucke et al., 2010; Kobayashi et al., 2017; Endicott and Brueckner, 2018; Xun et al., 2018). Given our finding of decreased acetylated ciliary tubulin, we were surprised to find neither the key acetylation enzyme αTAT1 nor the deacetylase HDAC6 to be significantly altered. Previous studies have shown that HDAC6 upon phosphorylation by Aurora A kinase (AurA) promotes tubulin deacetylation in cilia and ciliary disassembly (Pugacheva et al., 2007). This process can be prevented by BBS proteins through the recruitment of Inversin/NPHP2 to the ciliary base (Patnaik et al., 2019). However, AurA levels were not significantly changed, and phosphoproteomic (PP) analyses performed in parallel (Suppl. Fig. 3A/B; Suppl. Table 1) also showed no alteration in HDAC6 or AurA phosphorylation. Similarly, localization and expression levels of pHDAC6 in urine-derived BBS8^c.915delG^ URECs at the ciliary base was not altered (Suppl. Fig. 3C). However, our data revealed a significantly increased expression of another histone-deacetylase, HDAC2, in Bbs8^-/-^ kidneys. This finding could be further confirmed by Western blot experiments in both MEFs (Fig. 3D) and BBS8-deficient URECs (Fig. 3E).

**Figure 3.**
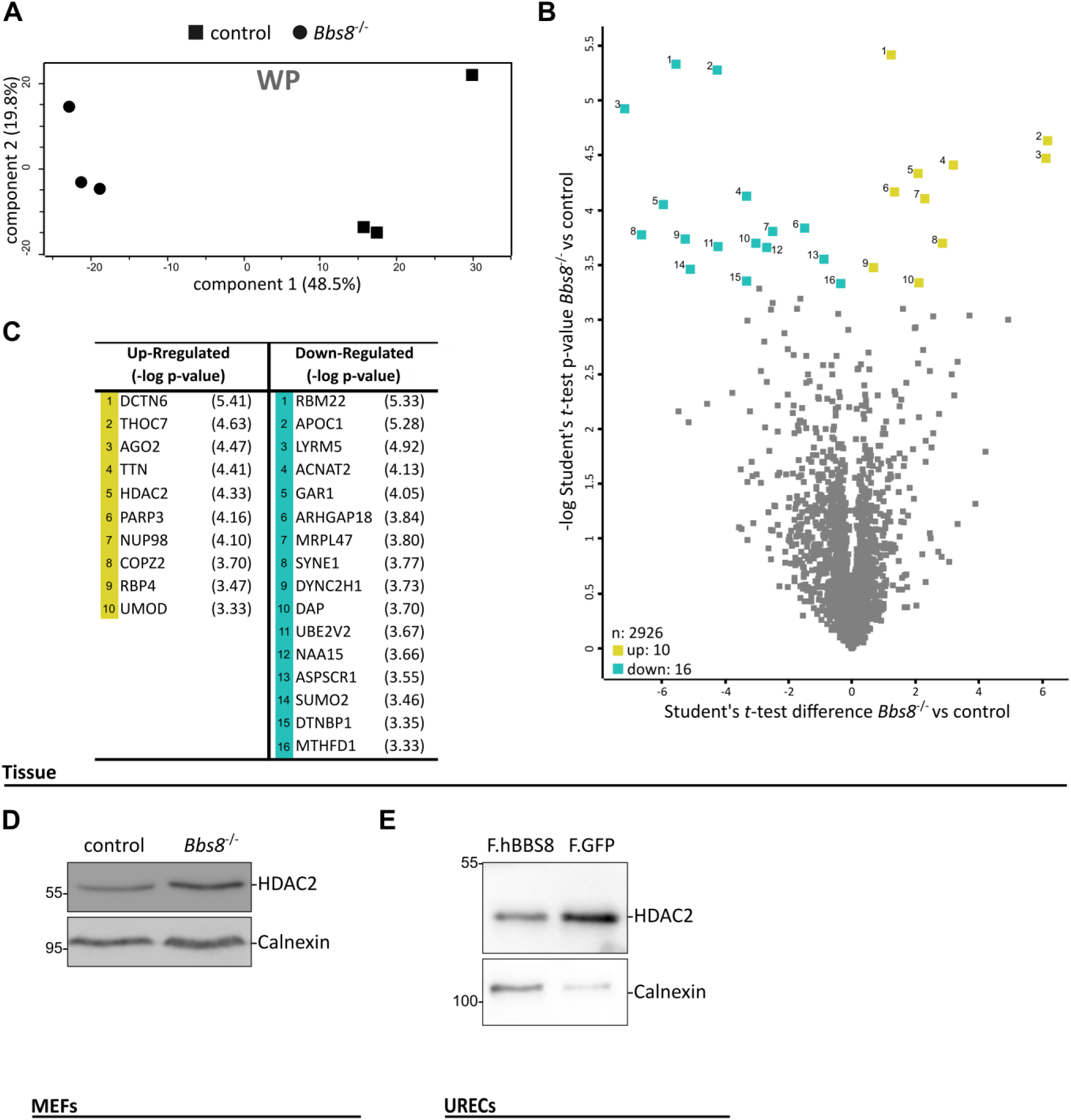
HDAC2 is one of the few proteins significantly upregulated in BBS8^-/-^ kidney tissue. (A) Principal component analysis (PCA) plot of the whole proteome (WP) of control and Bbs8^-/-^ kidney samples. The axes represent the percentages of variation explained by the principal components, three biological replicates per group. (B/C) Scatter blot of the proteome. Significantly (FDR≤0.05) up-regulated (yellow, 1-10) or downregulated (cyan, 1-16) proteins are marked and additionally listed in the table. (D) Lysates from control and Bbs8^-/-^ mouse embryonic fibroblasts (MEFs) were immunoblotted against HDAC2, with Calnexin as loading control. (E) Urine-derived BBS8^c.915delG^ URECs stably expressing either F.hBBS8 or F.GFP were immunoblotted against HDAC2 with Calnexin as a loading control.

### BBS8 co-precipitates with HDAC2

Intrigued by upregulation of HDAC2 upon loss of BBS8, we questioned whether there might be a connection between BBS8 and HDAC2 in the context of a shared protein complex. For an unbiased interactomic analysis, we exogenously expressed F.hBBS8 or an equally FLAG-tagged control protein in HEK293T cells and performed an anti-FLAG immunoprecipitation followed by MS/MS. The principal component analysis clearly separated the F.hBBS8 pulldown from the control (Fig. 4A). We identified 1151 proteins of which 986 were significantly enriched upon precipitation with BBS8 (Fig. 4B, Suppl. Table 2). Notably, among those were many known members of the BBSome (BBS1, BBS2, BBS4, BBS5, BBS7, BBS9) and the BBS chaperonin complex (CCT6A, CCT5, CCT3, CCT2, TCP1, CTT8, CCT4 and CCT7). We again found HDAC2 as the only enriched histone-deacetylase. Additional protein groups shown to potentially interact with BBS8 include heat shock binding proteins and proteins related to the Wnt signaling pathway (Supp. Fig. 4A). To confirm this potential interaction, we performed a similar co-immunoprecipitation experiment and were able to detect endogenous HDAC2 upon pull down with BBS8. (Fig. 4C). Co-immunoprecipitation in the reverse direction confirmed BBS8 co-precipitating with F9.HDAC2 (Fig. 4D).

**Figure 4.**
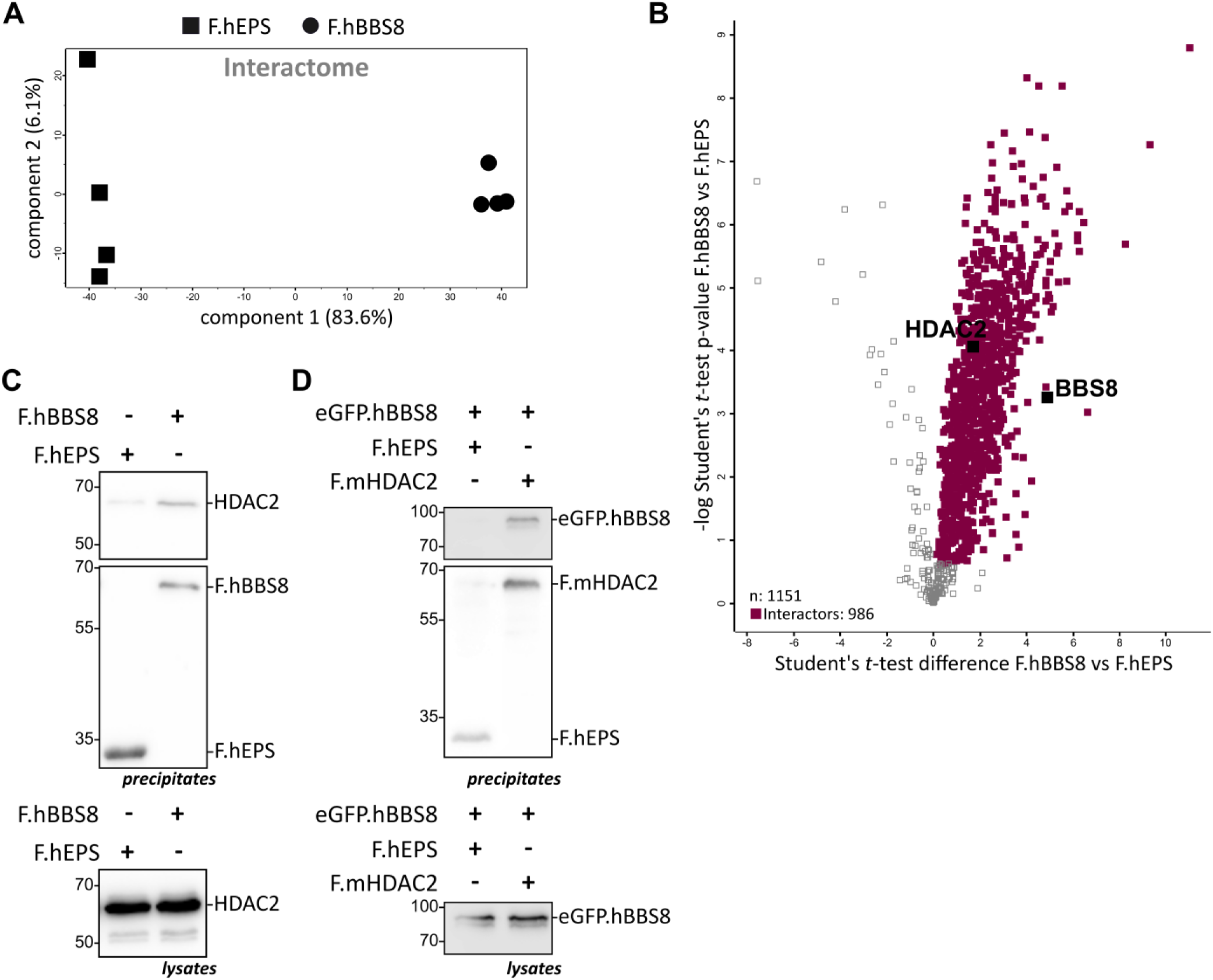
BBS8 and HDAC2 are components of a common protein complex. (A) Principal component analysis (PCA) plot of the interactome data of F.hEPS and F.hBBS8 in HEK293T cells. The axes represent the percentages of variation explained by the principal components. Four individual replicates were performed. (B) Scatter plot of the interactome, marked with potential interactors of BBS8 (FDR≤0.05) in magenta. (C) Co-IP from HEK293T cells, expressing F.hBBS8 or F.hEPS as control. Immuno-precipitation from lysates was done with α-FLAG (IP), immunoblot was performed with α-HDAC2 and with α-FLAG. (D) Co-IP from HEK293T cells expressing F.mHDAC2 or F.hEPS together with eGFP.hBBS8. Immuno-precipitation from lysates was performed with α-FLAG (IP), immunoblot with α-eGFP and α-FLAG.

### HDAC2 induces the deacetylation of alpha tubulin

HDAC2 has previously been associated with reduced cilia number in pancreatic ductal adenocarcinoma, a phenomenon attributed to increased transcription of AurA and consequently enhanced ciliary disassembly by HDAC6 (Kobayashi et al., 2017). However, our data from Bbs8^-/-^ tissue provided no evidence for elevated AurA expression or HDAC6 activity. Therefore, we aimed to demonstrate that HDAC2 directly influences tubulin acetylation. To this end, we transiently transfected HEK293 cells with F.mHDAC2 or F.GFP and included F.hHDAC6 as a positive control. Immunoblot analysis of cell lysates revealed that HDAC2 expression reduced of overall cellular levels of K40-acetylated α-tubulin, albeit to a slightly lesser extent than HDAC6 (Fig. 5A). This functional similarity between HDAC2 and HDAC6 is further supported by structural predictions using AlphaFold. The acetylated K40 residue of tubulin is predicted to interact with the HDAC domain of HDAC2 in a manner similar to its interaction with the second HDAC domain of HDAC6 (Suppl. Fig. 5). To further substantiate that the loss of ciliary tubulin acetylation in BBS8-deficient cells is attributable to HDAC2 activity, we employed the selective HDAC inhibitor Tucidinostat (TUC). TUC is a specific low nanomolar inhibitor of the class I HDAC enzymes (HDAC1/2/3; EC50: 100-200 nM) and class IIb HDAC enzymes (HDAC10; EC50: 100 nM) with almost no effect on HDAC6 (EC50 >10 µM) and has already been used in a clinical trial for patients relapsed or refractory peripheral T-cell lymphomas (Ho, Chan and Ganesan, 2020; Liu et al., 2021; Utsunomiya et al., 2022). We treated the urine-derived BBS8^c.915delG^ URECs expressing F.GFP or F.BBS8 with TUC (160 nM) or DMSO for 24 h in serum-free medium and stained for primary cilia with antibodies directed against ARL13B and acetylated-tubulin. As before, we observed in the control group (DMSO) a significantly decreased percentage of acetylated-tubulin positive cilia in the urine-derived BBS8^c.915delG^ URECS as compared to BBS8 re-expression. However, HDAC2-inhibition significantly increased the percentage of acetylated-tubulin positive cilia in BBS8^c.915delG^ cells, to a similar level as BBS8 re-expression did (Fig. 5B).

**Figure 5.**
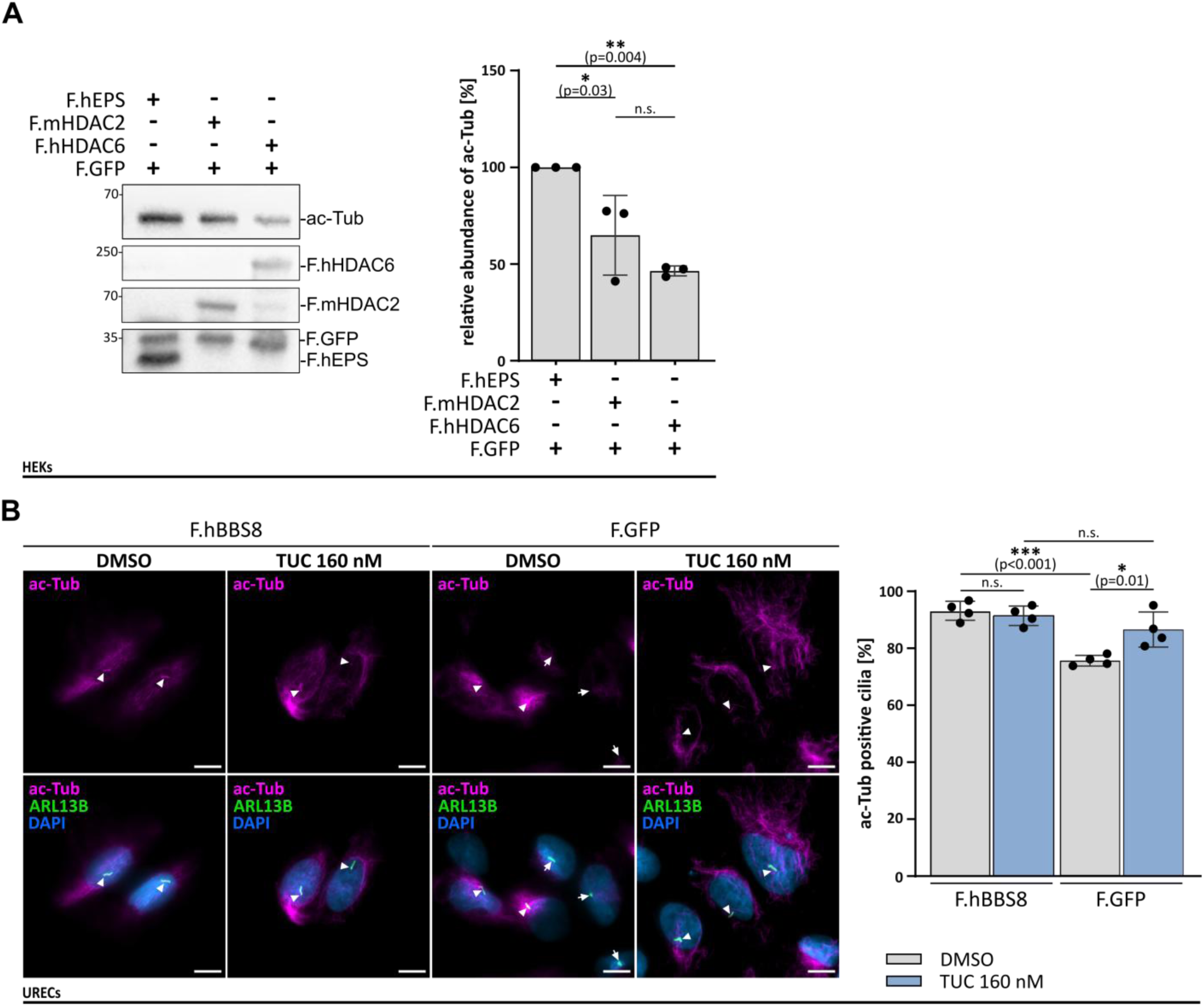
HDAC2 promotes tubulin deacetylation, and its inhibition increases tubulin acetylation in BBS8-deficient cells. (A) Lysates of HEK293T cells transfected with F.hEPS, F.mHDAC2, F.hHDAC6 and co-transfected with F.GFP were immunoblotted with α-acetylated tubulin and α-FLAG. F.GFP expression was used as control for quantitative comparison (n=3) with the statistical analysis of a one-way ANOVA followed by a two-sided Student’s t-test (p-value: >0.001***; 0.002**; 0.033*; ns1=10.12). Data were normalized to the acetylated tubulin amount in F.hEPS expressing cells. (B) BBS8^c.915delG^ patient-isolated URECs stably (re)expressing either F.hBBS8 or F.GFP were treated with the HDAC2 inhibitor Tucidinostat (TUC, 160 nM) or DMSO as control. Representative images of primary cilia co-stained with the cilia marker ARL13B (green) and acetylated-tubulin (ac-Tub, magenta; scale bar: 10 µm) displaying the restoration of acetylation within primary cilia in F.GFP BBS8^c.915delG^ URECs upon HDAC2 inhibition (n=4; arrowheads: ac-Tub positive cilia; arrows: ac-Tub negative cilia); graph showing the percentage of acetylated-tubulin positive cilia of the total amount of cilia determined by ARL13B staining. Statistical analysis was performed by using a one-way ANOVA followed by a two-sided Student’s t-test (p-value: >0.001***; 0.002**; 0.033*; ns1=10.12).

In summary, we conclude that loss of BBS8 results in a pathological upregulation of HDAC2 activity, which targets acetylated α-tubulin within the cilium, thereby perturbing ciliary dynamics and stability. Consistent with the relatively mild and late-onset kidney phenotype, these changes do not result in major alterations of cilia and have no impact on cilia length or number. Nevertheless, these subtle modifications in primary cilia architecture may affect ciliary dynamics and, over time, negatively influence the homeostasis of renal tissue. Thus, HDAC2 may indeed represent a promising therapeutic target. Supporting this notion, a chemical screen in two zebrafish models of polycystic kidney disease (PKD) identified both a pan-HDAC inhibitor and a class I HDAC inhibitor as suppressors of cyst formation (Cao et al., 2009). Our study provides a mechanistic basis for this positive effect of HDAC inhibitors by identifying the HDAC2-induced acetylation-defect of cilia which can be reversed by a class 1 inhibitor as potential driver of cystogenesis in BBS8, linked to subtle ciliary defects.

## Data Availability

The mass spectrometry proteomics data have been deposited to the ProteomeXchange Consortium via the PRIDE (Perez-Riverol et al., 2024) partner repository with the dataset identifier PXD064988 (interactome) and PXD064990 (proteomeBBS8-/- kidneys). Login details are available from the authors upon request.

## Supporting information

Suppl. Table 1: Proteome / Phosphoproteome Kidneys

Suppl. Table 2: Interactome

## Acknowledgements

We would like to express our sincere gratitude to Angelika Köser, Serena Greco-Torres, and Stefanie Keller for their exceptional technical assistance. We also deeply appreciate the outstanding support provided by the CECAD Imaging Facility and the CECAD Proteomics Facility. Our heartfelt thanks go to all members of our research group. Lastly, we extend our appreciation to the patient and their advocate for their generous contributions and commitment to advancing research. We would also like to thank Edward Seto for kindly providing us with the HDAC2 Plasmid. Molecular graphics and analyses were performed with UCSF ChimeraX, developed by the Resource for Biocomputing, Visualization, and Informatics at the University of California, San Francisco, with support from National Institutes of Health R01-GM129325 and the Office of Cyber Infrastructure and Computational Biology, National Institute of Allergy and Infectious Diseases (Meng et al., 2023). Funding: This study was supported by the German Research Foundation (DFG; SFB1403, project number 414786233, A09 to B.S. and T.B.; and FOR5547 to HMS, DW, and BS, project number 503306912, under Germany’s Excellence Strategy – EXC2151 – Project-ID 390873048 (to D.W.), and WA 3382/8-1 – Project-ID 513767027 (to D.W.). The NEOCYST consortium (Network for Early Onset Cystic Kidney Diseases) supported the project, with funding provided by the German Ministry of Education and Research (BMBF) under grant agreement number 01GM1515D. This funding (to B.S., M.C.L. and M.C.) had no impact on the selection of the topic or the content of the position paper. B.S. and M.C.L.. were also supported by the European Union (TheraCil). P.M. was supported by the Studienstiftung des Deutschen Volkes. The CECAD proteomics facility was supported by the large instrument grant INST 1856/71-1 FUGG by the German Research Foundation (DFG Großgeräteantrag).

## Author contributions

EK, HMS and BS designed the study. EK and JG performed most of the experiments. PAM prepared the experiments for MEFs. LKE performed and analysed the BBS8 interactome MS/MS experiments. HMS, CK, DW, MC, MCL, TB provided materials and support for the interpretation of results. EK assembled the final figures. EK and BS wrote the original draft. All of the authors discussed the project and contributed to the final version of the manuscript.

---

**Supplementary Figure 1.**
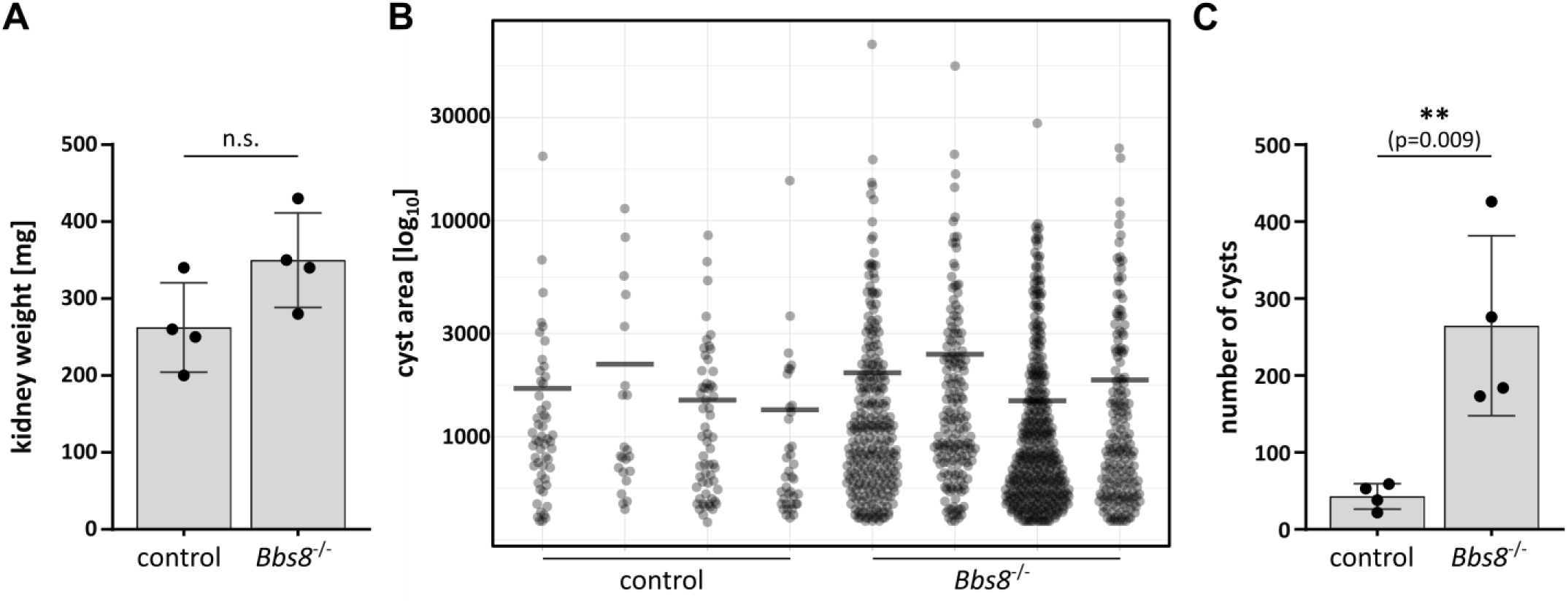
Small-cyst formation in Bbs8^-/-^ kidney and systemic inflammation. (A) Kidney weight of 46-week-old control and Bbs8^-/-^ mice did not significantly alter (n=4). (B) The measured cyst area is depicted as jittered dots with the mean as a horizontal line. The values are plotted on a log_10_ scale. Single individuals are plotted separately (n=4). (C) The overall number of cysts is significantly increased in Bbs8^-/-^ animals (n=4). For statistical analysis, a two-sided Student’s t-test was used (p-value: <0.001***; 0.002**; 0.033*; nslJ=lJ0.12).

**Supplementary Figure 2.**
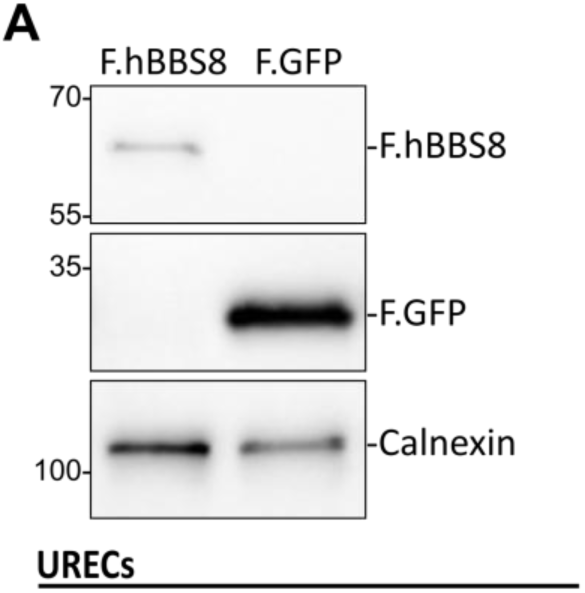
Re-expression of BBS8 in BBS8^c.915delG^ URECs. (A) Urine-derived BBS8^c.915delG^ URECS stably expressing either F.hBBS8 or F.GFP, were immunoblotted against FLAG with Calnexin as control.

**Supplementary Figure 3.**
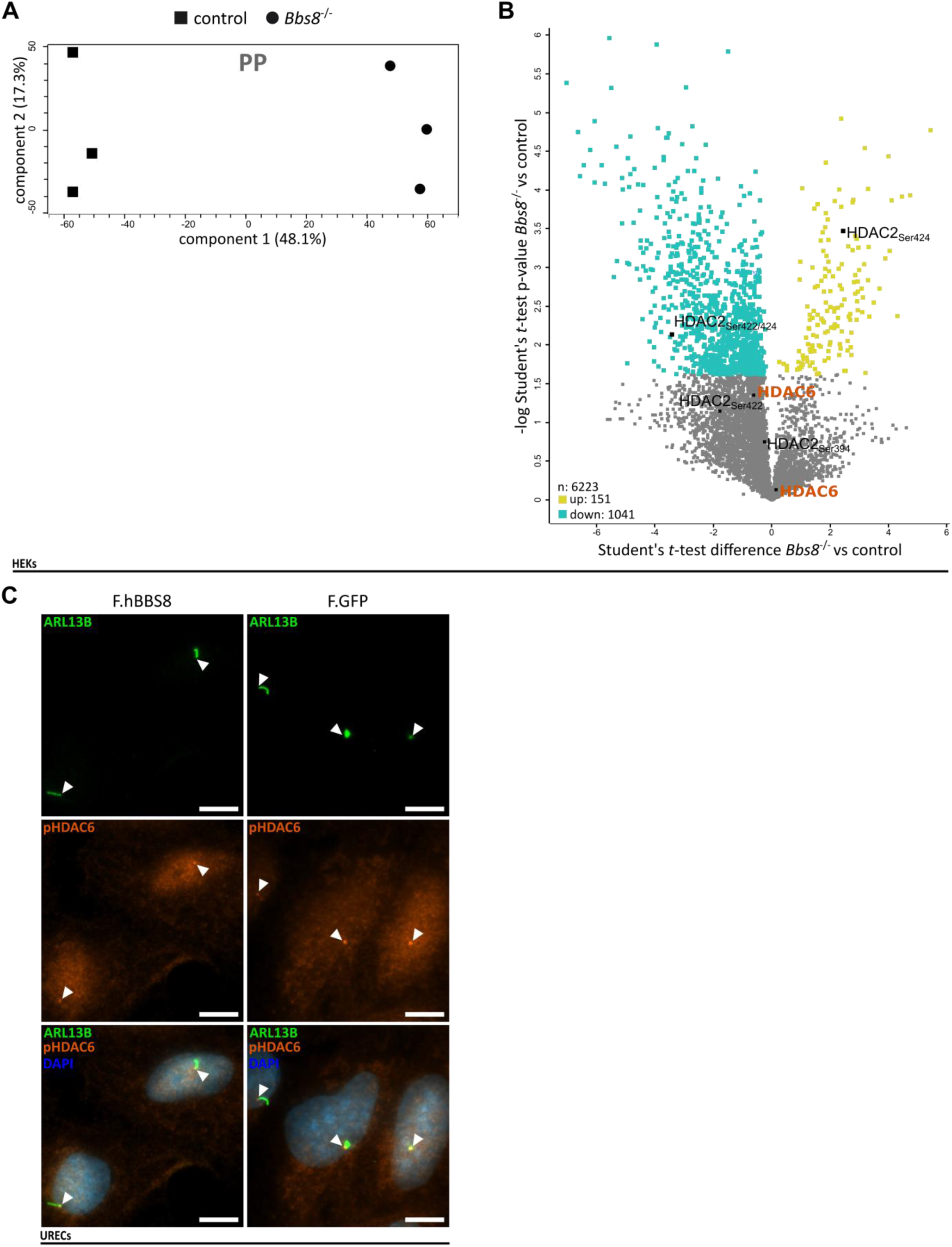
Additional analyses of the phosphoproteome data. (A) Principal component analysis (PCA) plot of the phosphoproteome (PP) data of control and Bbs8^-/-^ kidney samples. The axes represent the percentages of variation explained by the principal components. Three individual replicates were performed. (B) Analysis of the PP identified 6223 phosphosites, within a total of 2188 individual proteins from which 151 were significantly up- and 1041 were significantly down-regulated. Among these were multiple HDAC2 phospho-peptides containing 3 serine (Ser) residues: Ser394, Ser422 and Ser424. Notably, HDAC6 was not found to be changed in its phosphorylation. (C) Co-Staining for the cilia (ARL13B, green) and phopho-HDAC6 (orange) in urine-derived BBS8^c.915delG^ URECs re-expressing F.hBBS8 or F.GFP (Scale bar: 10 µm). Ciliary base is indicated by arrowheads.

**Supplementary Figure 4.**
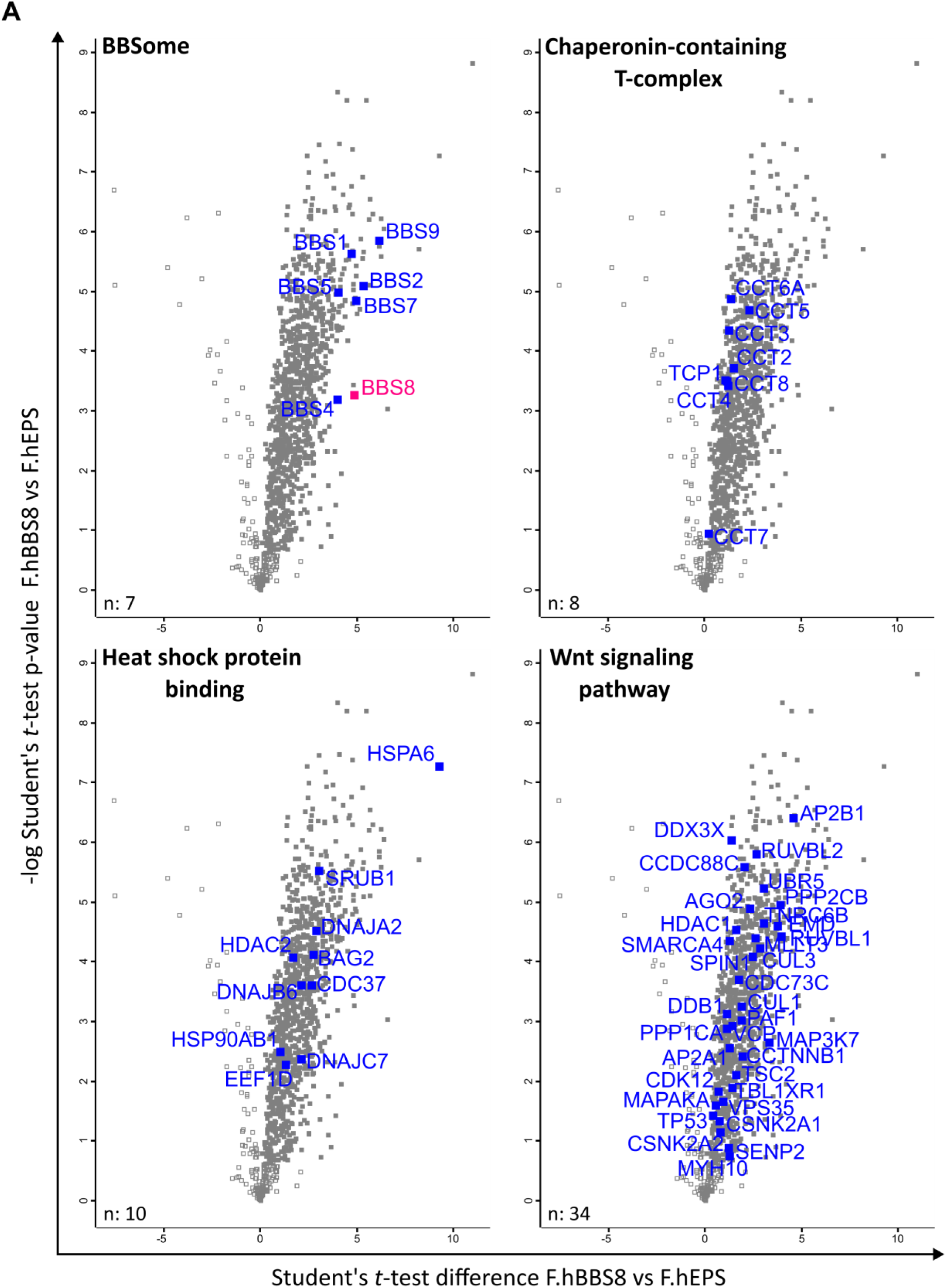
BBS8 interactors involved in cilia signaling pathways. (A) Scatter plot of different protein clusters expressed in the BBS8 interactome for different GO-terms: BBSome, chaperonin-containing T-complex, heat shock protein binding and Wnt signaling pathway.

**Supplementary Figure 5.**
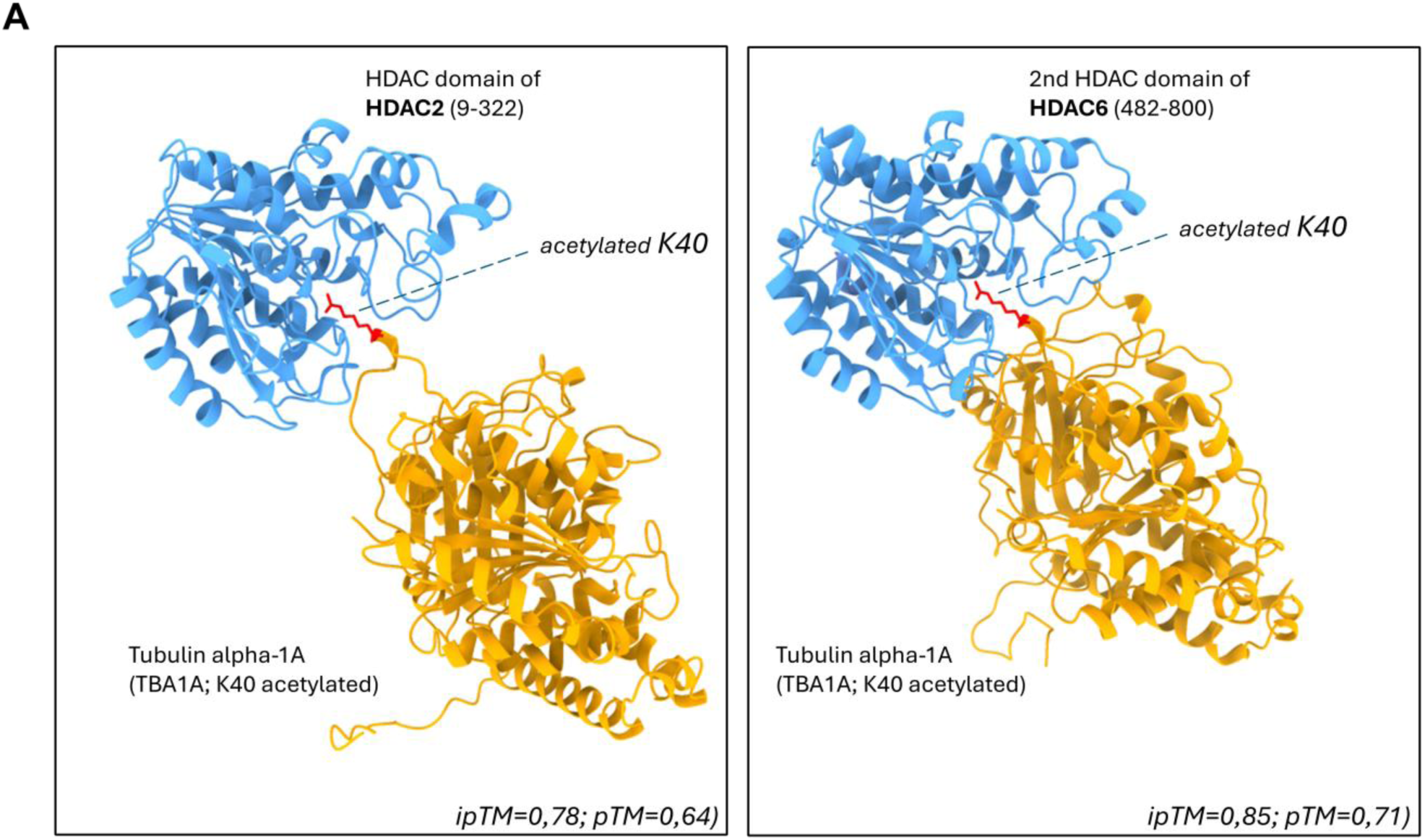
Modeling the interaction of HDAC2 and HDAC6 with K40-acetylated alpha Tubulin. (A) Human TBA1A (UniProt ID: Q71U36) with the PTM “N6-acetyl-L-lysine” added to K40 was co-predicted in AlphaFold with the HDAC domain of HDAC2 (Q92769; residues 9–322) or the second HDAC domain of HDAC6 (Q9UBN7; residues 482–800), a well-characterized tubulin deacetylase. The ipTM and pTM values were obtained from the AlphaFold server. The acetylated residue K40 is highlighted in red.

**Supplementary Figure 6.**
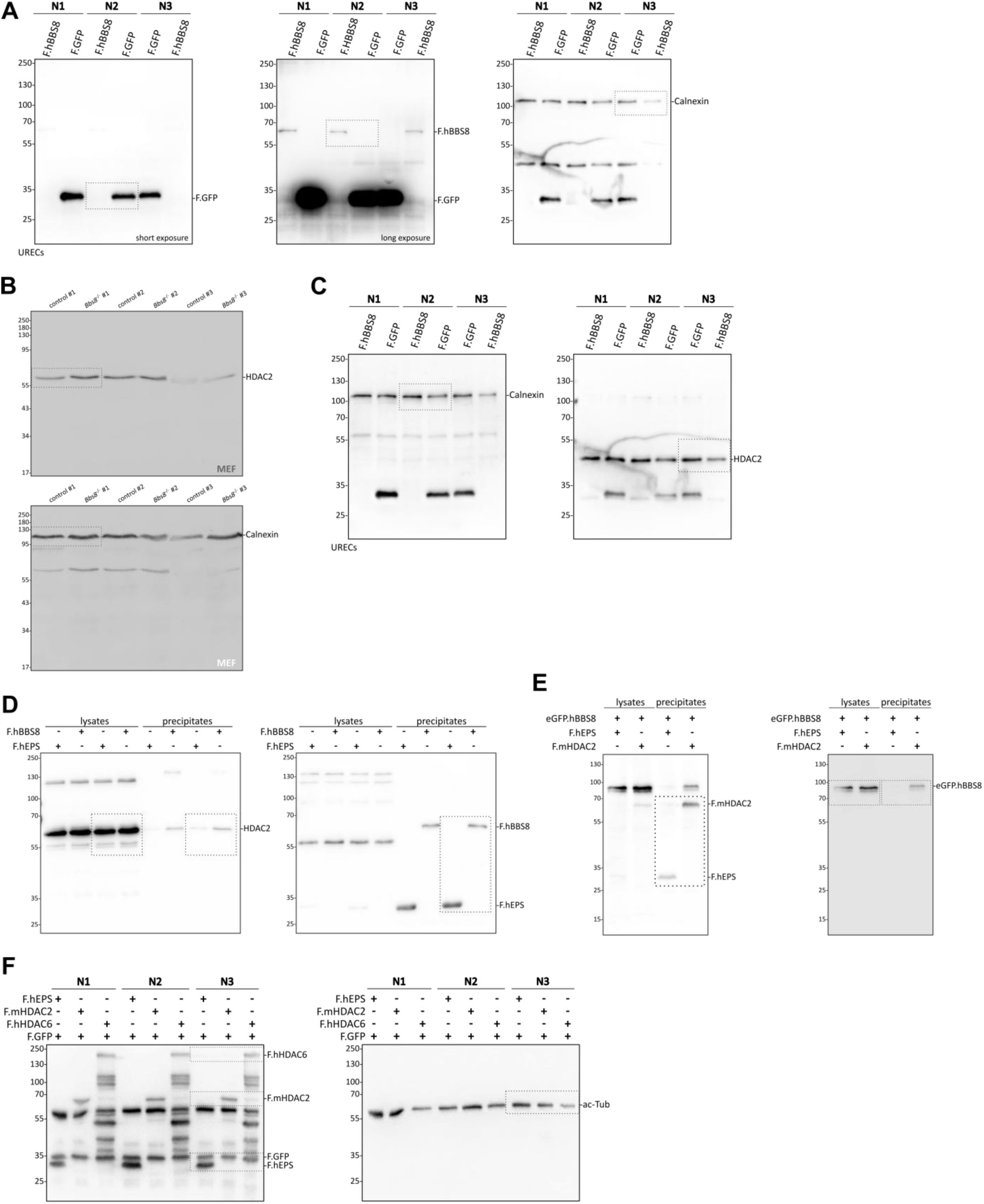
Original data: full-sized immunoblots. Original western blots only cropped to gel size (A) of Supp. Figure 2 A, (B) of Figure 3 D, (C) of Figure 3 E, (D) of Figure 4 C, (E) of Figure 4 D and (F) of Figure 5 A. To visualize the expression of F.hBBS8, the membrane was captured with a short and longer exposure time.

